# The unexpected consequences of predictor error in ecological model selection

**DOI:** 10.1101/2023.12.04.569928

**Authors:** Georg Manthey, Miriam Liedvogel, Birgen Haest, Michael Manthey, Joe Wynn

**Affiliations:** Institut für Vogelforschung “Vogelwarte Helgoland”, An Der Vogelwarte 21, Wilhelmshaven, 26382, Germany; MPRG Behavioural Genomics, Max Planck Institute for Evolutionary Biology, Plön, 24306, Germany; Swiss Ornithological Institute, Sempach, Seerose 1, Sempach, 6204, Switzerland; Institute of Botany and Landscape Ecology, University of Greifswald, Soldmannstraße 15, Greifswald, 17487, Germany

## Abstract

1. The ability to select statistical models based on how well they fit an empirical dataset is a central tenet of modern bioscience. How well this works, though, depends on how goodness-of-fit is measured. Likelihood and its derivatives (e.g. AIC) are popular and powerful tools when measuring goodness-of-fit, though inherently make assumptions about the data. One such assumption is absence of error on the x-axis (i.e. no error in the predictor). This, however, is often not correct and deviations from this assumption are often hard (or impossible) to measure.
2. Here, we show that, when predictor error is present, goodness-of-fit as perceived using likelihood will increase with decreases in sample size, effect size, predictor error and predictor variance. This results in predictors with increased effect size, predictor variance or predictor error being punished. As a consequence, we suggest that larger effect sizes are biased against in likelihood-based model comparison. Of note: (i) this problem is exacerbated in datasets with larger samples sizes and a broader range of predictor values - typically considered desirable biological data collection; and (ii) the magnitude of this effect is non-trivial given that ‘proxy error’ (caused by using correlates of a predictor rather than the predictor itself) can lead to unexpectedly high amounts of error.
3. We investigate the effects of our findings in an empirical dataset of wood anemone (*Anemone nemorosa*) first flowering date regressed against temperature. Our results show that the proxy error caused by using air temperature rather than ground temperature results in a ∆AIC of around 3. We also demonstrate potential consequences for model selection procedures with autocorrelation (e.g. ‘sliding window’ approaches). Via simulation we show that in the presence of predictor error AIC will favour autocorrelated, lower effect size predictors (such as those found on the edges of predictive windows), rather than the *a priori* specified ‘true’ window.
4. Our results suggest significant and far-reaching implications for biological inference with model selection for much of today’s ecology using observational data under non-experimental conditions. We assert that no obvious, globally-applicable solution to this problem exists; and propose that quantifying predictor error is key in accurate ecological model selection going forward.

## 2 Introduction

Model selection – the ability to rank statistical models based on how well they fit underlying data – is a central tenet of modern scientific statistics. Whilst early scientific endeavours focused primarily on experimental hypothesis testing, with the aim to distinguish between an *a priori* defined hypothesis and an alternative null hypothesis (Popper, 1935), modern ecology is increasingly independent of this paradigm for several reasons. First, ecological systems are often ‘large and slow moving’, making hypotheses hard to test experimentally (Wolkovich *et al*., 2012; Tredennick *et al*., 2021). This has lead to a focus on correlative analysis using ‘natural’ experiments (e.g. (Wynn *et al*., 2020; Acker *et al*., 2021)), which often requires variable selection. Second, the complexity implicit in ecological systems means that ecological problems are often approached with limited knowledge on how and when a predictor might interact with a response variable. One solution to this problem that has emerged (e.g. Gienapp *et al*., 2005; Kruuk *et al*., 2015; Wynn *et al*., 2022) is to test many different predictors - or repeatedly test different versions of the same predictor - and compare goodness-of-fit to generate new scientific hypothesis (Tredennick *et al*., 2021).

This hyper-exploratory ecological approach has marched in lockstep with increasing computing power, machine learning and ever-more complex and nuanced parametric statistical techniques (e.g. Bates *et al*., 2009; Payo-Payo *et al*., 2022; Schirmer *et al*., 2023), and has revolutionised environmental science. By allowing biologists to test multiple hypotheses, model selection has allowed us to discover trends that are, from the outside, extremely difficult to identify *a priori*. For example, in systems where year-on-year survival changes through time - but is driven by an as-yet-unknown causative agent, with an as-yet-unknown temporal lag - model selection can be used to identify the environmental cause (e.g. the effect of extreme weather on seabird mortality; Frederiksen *et al*., 2008; Genovart *et al*., 2013). However, model selection techniques are necessarily only as good as the goodness-of-fit metrics ascribed to them, and hence effective model selection is contingent upon unbiased and accurate characterisation of model fit in all instances.

One of the simplest measures of goodness-of-fit is the r^2^ (‘r-squared’) of a given model, which is defined as the proportion of the variance for a response variable that is explained by one (or more) predictor variables. However, r^2^ is necessarily inflated by the inclusion of multiple predictors and, in any case, can only be fitted to a select number of statistical models. It is, therefore, of limited utility in biological model selection. The use of likelihood functions presents an extremely powerful (and in principle ubiquitous) alternative, with likelihood-based metrics such as the Akaike Information Criterion (‘AIC’), the Bayesian Information Criterion (‘BIC’) or likelihood ratio testing being perhaps the most popular likelihood-derived measure of goodness-of-fit (Akaike, 1976; Schwarz, 1978; Foster & George, 1994). AIC, in particular, is extremely popular owing to its generalisability across statistical contexts, and is itself not susceptible to the same parameter inflation as r^2^ (Burnham & Anderson, 1998).

As with all parametric statistics, likelihood-based measures of goodness-of-fit are based on several assumptions about the data in order to function. Whilst many of these assumptions are well known in the biological sciences (e.g. that the data adheres to the distribution stated in the underlying model), the assumption of a perfectly measured predictor variable is perhaps not as widely known. Indeed, biologists tend to optimise the accuracy of their response variables, with perhaps less attention paid to measuring predictors as accurately as possible. This is particularly true when considering climatic predictor variables, which are often remotely-sensed and hence could deviate from local conditions in ways that are very hard to predict or measure (e.g. Thorup *et al*., 2017; Aikens *et al*., 2020; Kempton *et al*., 2022).

Here, we investigate the effect that predictor error might have on biological model selection using likelihood and AIC. Specifically, we attempt to determine a) whether a theoretical basis exists that leads us to question the use of likelihood and its derivatives in the presence of predictor error; b) whether this theoretical basis has any biological relevance in a real-world scenario; and c) whether any problems encountered can be solved. This we did by deriving the gaussian likelihood function to include predictor error, and then using simulation and real-world data to validate and assess the impact of our findings.

## 3 Methods

### 3.1 Simulations and derivations

All simulations and visualisations were conducted in Python 3.10 (Python Core Team, 2021) using the libraries numpy (Harris *et al*., 2020), pandas (McKinney, 2010), matplotlib (Hunter, 2007) and ipython (Pérez & Granger, 2007). We used the formula for a general error-in-variables linear regression (Equations 1 and 2; Fuller, 1987), and from this sought to derive the effect of predictor error on log-likelihood given a least-squares method not accounting for predictor error. From this we calculate the effect of predictor error on AIC. This derivation is shown in the results section.

We used our derivation of likelihood in the presence of predictor error to investigate the effect of temporal and spatial autocorrelation on model selection. This we did by calculating the effect of predictor error on likelihood for a model where effect size was autocorrelated across parameterspace, with the highest effect sizes seen at the point in space representing the ‘true’ predictive window and decreasing with increasing distance from this window.

### 3.2 Real-world example

To investigate the implications of our theoretical framework we wanted to quantify predictor error in a real-world biological system. Given how extremely context-specific the implications of our findings are likely to be, this real-world implementation should not be considered a generalisable rule, but rather a proof-of-concept that might allow others to investigate errors more cautiously in their own systems.

Our proof-of-concept was parameterised using data from the Climate Data Center OpenData of the Deutscher Wetterdienst (German weather service; ‘DWD’; Kaspar *et al*., 2013). Specifically, we looked at the effect of predictor error in a system where error is implicit, which we did by considering the effect of air temperature on the first flowering dates of wood anemones (*Anemone nemorosa*; see Figure 3b). The wood anemone flowers within 2 weeks of first sprouting, with the flower being fully developed in the previous autumn (Shirreffs, 1985). The first flowering date must, therefore, be causally closer related to ground temperature than air temperature since the plant does not experience air temperature until extremely close to the point of flowering. Both air and ground temperature are measured at weather stations across Germany, and wood anemone first flowering dates are also observed and recorded at some of these weather stations. Hence, this seems to represent an ideal system allowing the investigation of the effect of predictor error on likelihood-derived goodness-of-fit.

**Figure 1.**
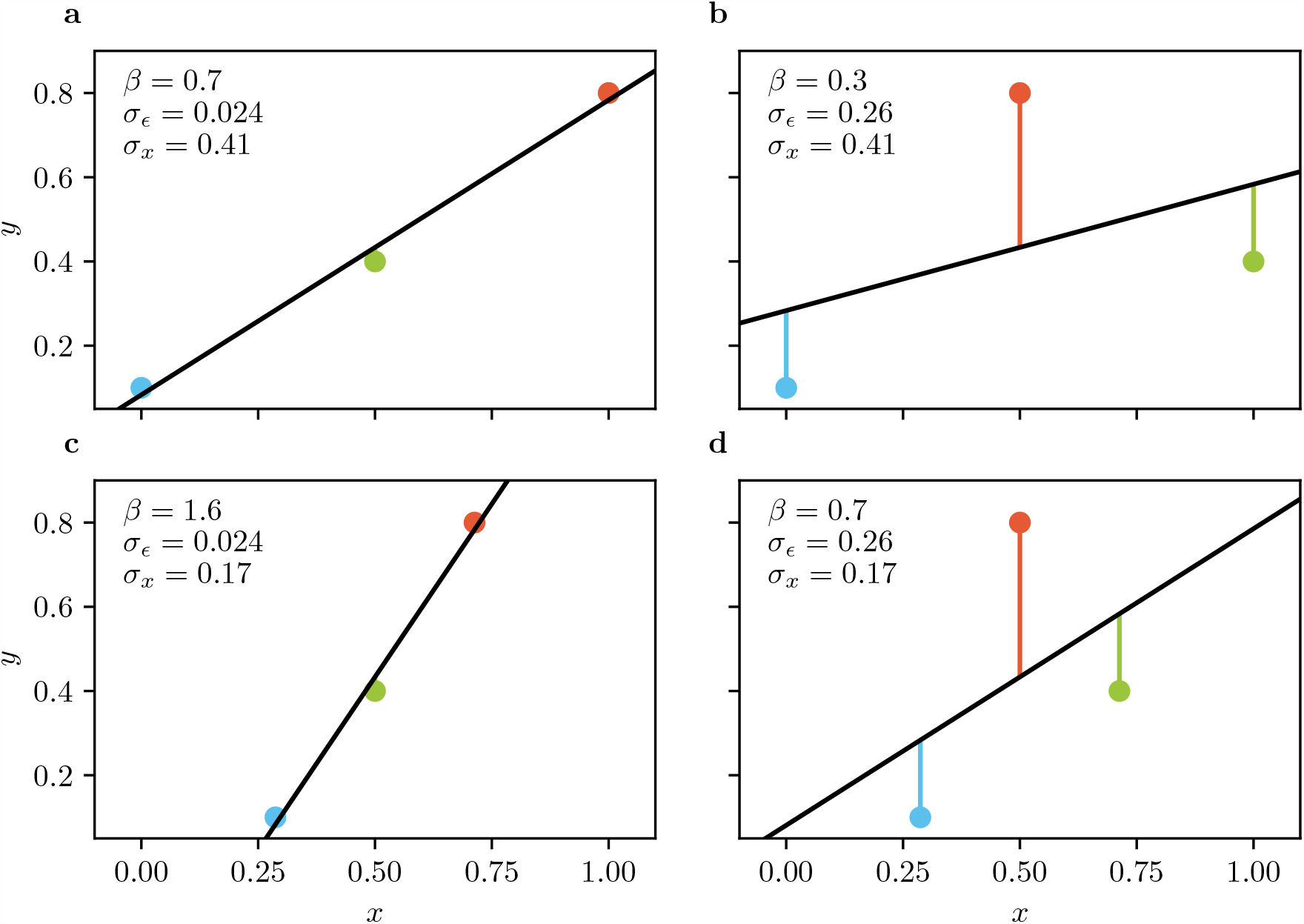
The relationship between variance and gradient in model selection. A visualisation of Equation 8. **a)** A simple linear regression through 3 points (blue, green, red) with low residual variance (*σ*_*ϵ*_), high effect size (*β*) and high predictor variance (*σ*_*x*_). **b)** By swapping corresponding values in the predictor (*x*, blue, red, green), the effect size can be decreased. At the same time, this increases the residual variance, while predictor variance stays constant. **c)** The effect size can also be altered by varying the predictor variance, while keeping residual variance constant compared to **a). d)** Reducing the predictor variance may also increase the residual variance, here while keeping the effect size constant compared to **a)**. All 4 plots shown share a common response variable (*y*) with values 0.1, 0.4 and 0.8.

**Figure 2.**
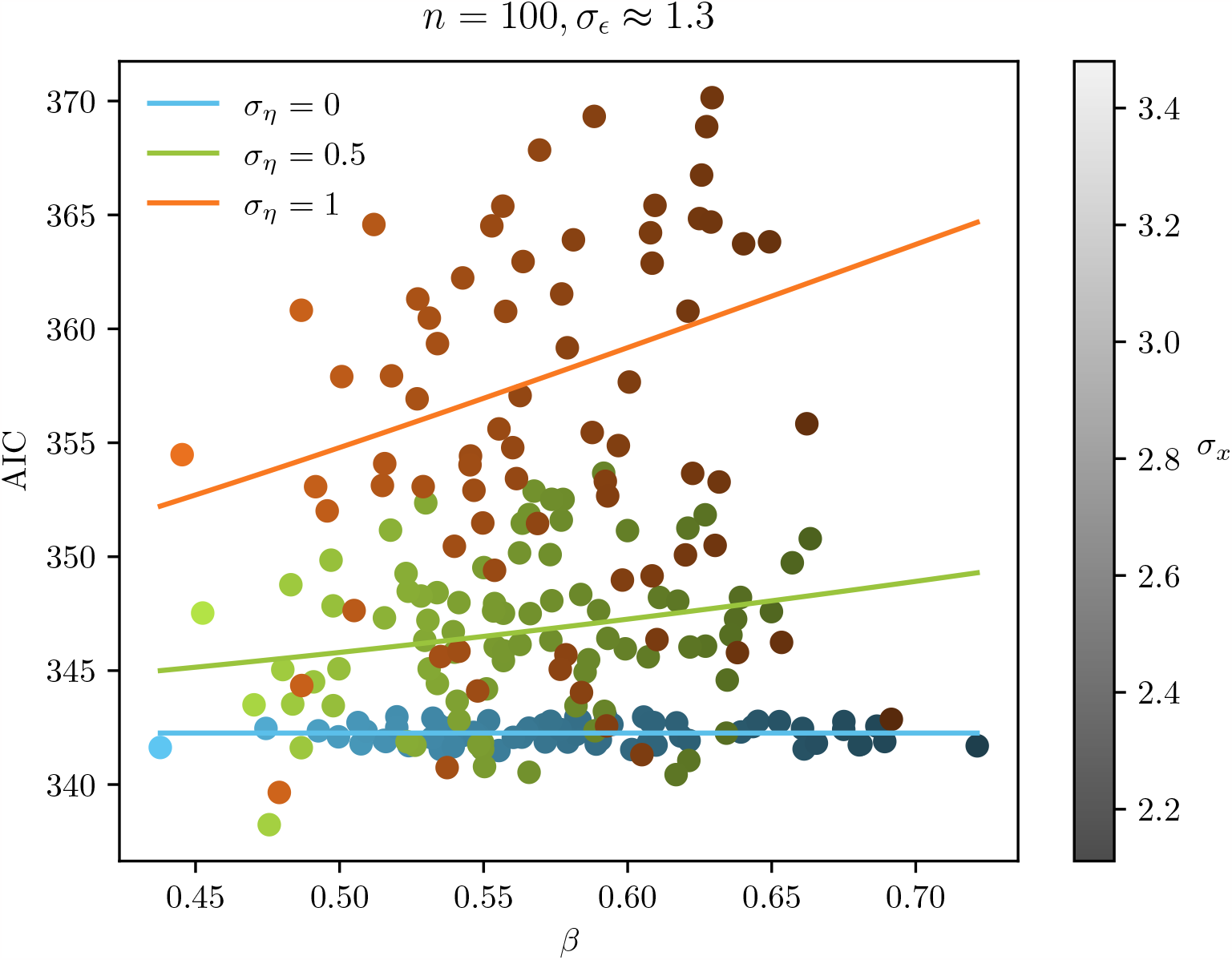
Visualisations of Equation 7, showing that when predictor error is present, AIC is correlated with gradient (*β*), predictor variance (*σ*_*x*_, shading) and predictor error (*σ*_*η*_, blue, green, red). The coloured lines show the ideal AIC value for the given parameters according to Equation 7, which are backed up by simulation results shown by the scatterplot. This also shows that effect size and predictor variance interact (see Box 1), as shading correlates with *β*. This correlation also leads to a perceived opposite effect of *σ*_*x*_ on AIC compared to what Equation 7 predicts. Every simulation result shown uses the same response variable and has about the same residual variance (and should, therefore, have the same AIC) before adding predictor error.

**Figure 3.**
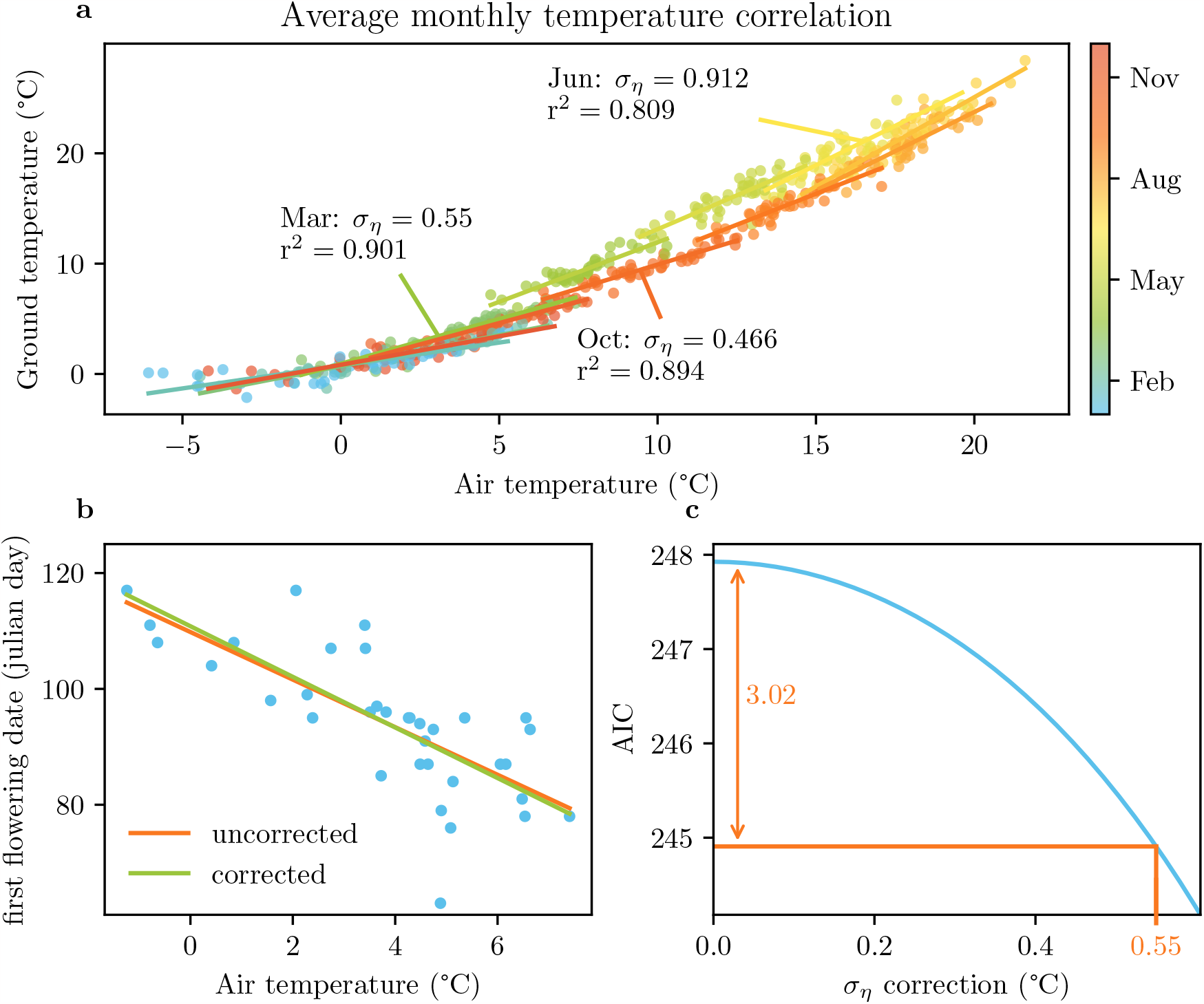
A case study to illustrate the effect of predictor error on AIC. **a)** Regression of air temperature against ground temperature for each month over 35 years at a weather station in Greifswald, Northern Germany. The error in this relationship can be considered the ‘proxy error’ (in march *σ*_*η*_ = 0.55 °C) associated with using one variable as proxy, when the other would be more appropriate. **b)** The regression of air temperature against the first flowering date of wood anemone at the same study site in Greifswald (blue dots). Regression without correcting for the predictor error associated with using air temperature instead of ground temperature is plotted in orange, in green the corrected regression. **c** Illustrates the change in AIC for different amounts of estimated predictor error (blue curve). The error extracted from the air temperature to ground temperature correlation (*σ*_*η*_ = 0.55 °C, orange) would yield an AIC difference of 3.02.

Daily temperature values for air (2m above ground) and ground (5cm below ground) were downloaded for a weather station in Greifswald, North-East Germany (54.0967°N,13.4056°E), from which a mean monthly ground and air temperature was derived. Given the time of year in which first flowering is typically recorded, March temperatures were used in a linear regression against first flowering date. This regression was used as our case study for investigating the effect of predictor error on likelihood-based measures of goodness-of-fit, with ground temperature assumed to be the ‘true’ predictor to which air temperature could act as a proxy. The variance in this relationship was, therefore, assumed to be a reasonable estimation of a system containing error.

Because both ground temperature and air temperature are measured at the same point in space, but at a different (though nonetheless close by) point to the plants from which flowering dates are recorded (since the weather station cannot be ‘on top of’ the plant), the exact coefficients of the relationship presented here may vary slightly to the ‘true’ value. Nonetheless, these effects are likely to be very small, and hence we believe that the relationship between air temperature and ground temperature is a suitable model for the effects of predictor error in biological systems.

## 4 Results

### 4.1 Deriving likelihood to include predictor error

We focus on a simple linear model to investigate the effect of a variable *x* on a variable *y* with effect size *β*, intercept *α* and response error *ϵ* (mean 0 and variance 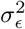; Gauss, 1809; Fuller, 1987):

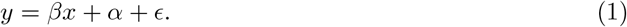

In addition to the standard model shown above we can also consider error in the predictor *η* (mean 0 and variance 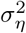) as well as our focal predictor *x*, leading to the observed predictor used in regressions going forwards (*w*) being defined after Fuller (1987):

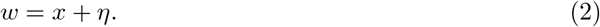

It is widely recognised that this kind of error in the predictor leads to regression dilution, which is defined as shallower estimated gradients with increased predictor error even when effect size is held constant by the following Equation: (Spearman, 1904; Frost & Thompson, 2000):

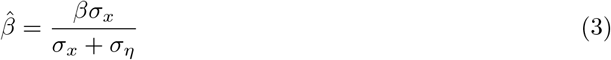

We can, however, use this model of predictor error to also infer how predictor error interacts with common measures of goodness-of-fit by including predictor error in the formula for (log)-likelihood:

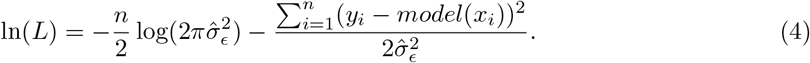

In this equation, we note that the only term affected by predictor error is the estimate of model error variance 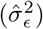. When there is error present in the predictor, the use of a classical maximum likelihood estimator to infer this value yields the following over-approximation (see SI derivation 2):

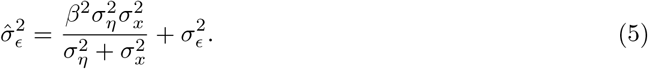

Perhaps unsurprisingly, the estimator accounts for the additional error in the system by incorporating it into the estimation of 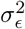. This is akin to error in the predictor being assumed to be error in the response. However, the interaction with the predictor variance 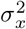 and the effect size *β* is unexpected (see Figure 2).

When restricting ourselves to realistic variable combinations that could occur during model selection - *β* (effect size), 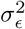 (response variance), *σ*_*x*_ (predictor variance) and *σ*_*y*_ (response variance; together we term these variables ‘meta-variables’) co-vary, which restricts parameterspace (see Box 1) - we can integrate Equation 8 from Box 1 to remove the dependence of one of the meta-variables. Removing for example the dependence on *β* in Equation 5 yields the following over-approximation (see SI derivation 3):

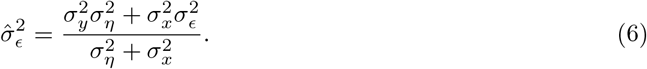

This equation could, at least in theory, be further simplified, by normalizing predictor and response (yielding 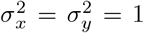). However, due to correlation between the meta-variables outlined in Box 1, a change in the predictor and response variances would also yield a changed residual variance, resulting in exactly the same overestimation. This is, then, of little utility and does not affect the effect of predictor error on goodness-of-fit as estimated via likelihood.

We can combine Equations 4 and 6 to estimate the effect of predictor error on likelihood, which yields the following equation:

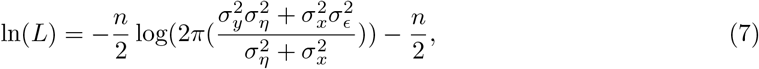

suggesting a negative correlation of log-likelihood with 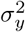 (response variance), 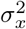 as well as 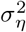. As information criteria such as AIC are perfectly correlated with log-likelihood (for a given number of parameters (*k*)), they will vary identically, as can be seen in Figure 2. In turn, this suggests that goodness-of-fit, as inferred via AIC or similar metrics, will be *lower* in cases of higher effect sizes, and that this effect is more pronounced when predictor error or variance are *higher*. This means that likelihood, AIC and other likelihood derviatives will scale with factors entirely unrelated to goodness-of-fit.

### 4.2 How large do we expect predictor error to be in a real-world example?

Quantifying predictor error is difficult, since the sources of predictor error are typically unknown. Measurement errors are more easily quantifiable and reducible by performing multiple (independent) measurements and averaging over these measurements, though error caused by device inaccuracy is unlikely to be the only source of error in biological modeling. We suggest that error induced by indirectly measuring predictor variables - which we call ‘proxy error’ - represents a non-trivial contribution to error overall.

In ecology, remotely-sensed data (e.g. from weather re-analysis or satellite observation) - used as a proxy when more accurately measured local information is not available - has proven an invaluable addition to the model selection framework. However, any deviations from an exact causative relationship between two biological events will necessarily induce predictor error. For example, when considering the response of a top predator (e.g. a pelagic seabird) to a changing climate (e.g. an increase in sea surface temperature), each causative level between the predictor and the response (e.g. fish reproductive success) will introduce proxy error if the correlations are anything other than perfect (e.g. Fort *et al*., 2012). The same can also be true in more direct causal instances; even if the flowering of a tree, for example, was directly regulated by temperature, then using remotely-sensed data for an entire forest is likely to be unrepresentative of the local conditions encountered by an individual tree and hence error might be induced (e.g. Büntgen *et al*., 2022).

Given our above derivation of likelihood with predictor error, we suggest that these errors could have a measurable effect on goodness-of-fit. However, exploring how such errors affect measures of goodness-of-fit in real-life scenarios is extremely difficult. That said, through the comparison of different data sources - that are either more or less proximal to the event in question - we might obtain a minimum value for the proxy error associated with a regression and, hence, calculate a minimum change in perceived goodness-of-fit.

To estimate this, we quantified error in a real biological system. For this we chose the effect of air temperature on the first flowering date (see Methods) of the wood anenome at a site in north-east Germany. Linear regression of air temperature analysis of first flowering date against mean March air temperature - with March chosen since it directly preceded the earliest first flowering date (Sparks *et al*., 2000; Tooke & Battey, 2010) - showed a highly significant correlation that explained a large amount of variation (F = 40.04, r^2^ = 0.548, p *<* 0.001, AIC = 249.9, Figure 3c). This is in spite of the biological causality most likely acting through the ground temperature, as outlined in the methods section.

To estimate the effect of this source of error on first flowering date, we parameterised the relationship between air temperature and ground temperature. Perhaps unsurprisingly, there is an extremely strong correlation between mean monthly air and ground temperature for temperatures above freezing (F = 383, p *<* 0.001, r^2^ = 0.901, AIC = 76.2, Figure 3a). However, whilst the two variables are clearly highly dependent (as evidenced by the extremely large r^2^), there is still some variability between the two. This can be expressed as the mean of the squared residuals (*σ*_*ϵ*_). In this case, *σ*_*ϵ*_ is 0.55, which can be assumed to be the value of proxy error associated with air temperature when used as a proxy for ground temperature. This value can be inserted into Equation 3, which in turn can be used to calculate a change in gradient (see Figure 3c).

Using Equation 7, we can therefore calculate a correction factor for log-likelihood and, in turn, AIC. This allows us to estimate the change in AIC associated with our original air temperature regression, should predictor error be eliminated. For this regression, we found that predictor error causes a change in AIC of 3.02 (see Figure 3d), a value that is slightly above the threshold typically considered ‘significant’ in model selection (Burnham & Anderson, 2001). That the ∆AIC is large enough to affect model selection is perhaps surprising, since the error coefficient underpinning this relationship was relatively small (0.55) and the two variables correlated near-perfectly (r^2^ = 0.901).

This would suggest, therefore, that even when predictor error is small it is sufficient to potentially disrupt common model selection protocols.

However, we note that the error coefficient observed between air and ground temperature varies month-by-month (see figure 3a), meaning that the amount of error in the system is not constant through time. Specifically, the air temperature/ground temperature relationship gives error coefficients that vary between 0.47 (October) and 0.934 (June). This means that the error in this relationship does not simply impose an intercept on AIC values, and instead adds different amounts of error for different months. In the case of June, this would lead to AIC improving by a value of 11.2 in the case of an underlying relationship akin to the wood anemone example outlined above, contrasting sharply with the ∆AIC of 2.21 observed in October. This makes the effects of predictor error still harder to quantify in biological systems, and may have serious implications for comparisons of the same variable measured at different times.

## 5 Discussion

Here, we demonstrate that surprisingly small amounts of error in a predictor variable can produce measurable and non-trivial changes in likelihood-based measures of goodness-of-fit (including popular metrics such as AIC; see Figure 3). This comes about through a multiplicative interaction between sample size, effect size and predictor variance that causes overestimation of response error (see Figure 2) which, in turn, leads to an underestimation of goodness-of-fit. Based on estimations of error derived from a real biological system, we believe these underestimations are non-trivial (see Figure 3). Crucially, we find that this effect is most pronounced in instances of greater effect sizes, sample sizes and predictor variance - all facets typically considered ‘desirable’ in ecological modeling. This, perhaps perversely, suggests that our findings are especially relevant to very large and well parameterised datasets.

### 5.1 How large a problem is this?

This effect of error on likelihood-based goodness-of-fit metrics is necessarily most important when considering model selection. If all variables are beholden to the same changes in estimated goodness-of-fit then there will, inevitably, be no relative change. However, given that predictor variables will have different variances and effect sizes this seems unlikely. Further, since predictor error is not necessarily well parameterised in all instances, knowing the extent of this problem is intrinsically challenging.

Whilst it is not our intention to make assertions about predictor error in specific instances, we nonetheless aim to advise on instances where this problem might be most severe. Quantifying the effect of our findings is inherently difficult, as there are very many instances where the source of predictor error is unclear and hence the estimation of its magnitude is difficult or impossible (e.g. in remotely-sensed variables with no available ground-truthing). Estimates of error in these remotely-sensed datasets often take the form of root mean squared error (RMSE) - deviations from a 1:1 ratio with some form of validation data (e.g. through comparison with a weather station) - and such comparisons suggest much larger errors than we demonstrate in our wood anemone example (e.g. Hersbach *et al*., 2020; Jiang *et al*., 2021; McNicholl *et al*., 2021). This could, therefore, suggest that such datasets might be surprisingly susceptible to AIC inflation. However, RMSE is an imperfect measurement of error in this context, since it does not consider that imperfect (i.e. not 1:1) correlations not necessarily lead to biases in likelihood estimation. This is because measurement error in this context is measured relative to the least-mean squares regression line, rather than an imposed 1:1 correlation (see Figure 3a). As such, whilst current estimates of error in remotely-sensed data seem substantial, extensive parameterisation of climatic models relative to real-world data is required to truly understand the extent of this problem. This is because the effect of predictor error would seem to be extremely context-specific, and very hard to predict *a priori*.

The other situation in which we suggest our findings may make model selection problematic is in multiple hypothesis testing of different versions of the same variable (e.g. comparing minimum, maximum and mean of temperature). Since the effect of predictor error is heightened in instances of increased effect size, model comparisons of the same variable with different effect sizes will cause smaller effect sizes to win out. This could either result from including multiple summary statistics derived from the same model, or through the inclusion of the same predictor sampled at different points across time/space (see below).

#### ‘Edges and echoes’: the special case of ‘sliding window’ model selection

The problem presented by multiple hypothesis testing with the same variable is particularly pronounced when considering spatially and/or temporally auto-correlated versions of the same variable - as is encountered in sliding window approaches to climatic model selection (e.g. Bailey and Van De Pol, 2016; van de Pol *et al*., 2016; Haest *et al*., 2018). In such instances, we might find that the edges of a climatic window (the centre of which has genuine predictive power) might out-compete the window itself. This is because the edge likely has a lower effect size, and hence will be less affected by predictor error. Alternatively, in the case of cyclic variables (e.g. annual variation in climate) we might expect autocorrelation between variables measured on the same date in different years, which in turn might cause biological effects to ‘echo’ between years in a model comparison, leading to the ‘wrong’ year’s climatic window being selected.

Whilst the above is undoubtedly true if effect size alone decreases with distance to the ‘true’ window, in reality effect size co-varies with (and is constrained by) by predictor variance and overall model error (see Box 1). This means that not all parameter space is necessarily possible, and in turn that we do not expect this ‘edges and echoes’ paradigm to play out in all instances. For example, if predictor variance and effect size were to act antagonistically, then model selection would always target the correct window. However, if this were untrue then we might expect it to miss.

Knowing whether a sliding window hits or risks missing its intended target is extremely difficult to know *a priori*, since predictor variance and effect size cannot be measured accurately due to predictor error (see Equation 3). This means that analysing this phenomena in detail is likely impossible. Nonetheless, we can construct a theoretical example using Equations 6 and 7 in which an autocorrelated, lesser fitting model will outcompete the ‘true’ model (see Figure 4). This suggests that the ‘edges and echoes’ effect is possible in sliding window model selection. However, by varying the input parameters of our simulation, situations where this is not the case can also be produced. This would suggest that sliding-window approaches to model selection can miss their intended targets, but do so in a manner that is almost impossible to predict. This paradigm becomes still more complex when we consider that the error between two variables is not constant through time. As noted in our worked example, the error in a given system can vary (see Figure 3), meaning that the effect of predictor error on goodness-of-fit becomes even harder to estimate. This makes the reliability of AIC-based model selection even harder to predict, and as such we might suggest that sliding-window model selection should be approached with caution.

**Figure 4.**
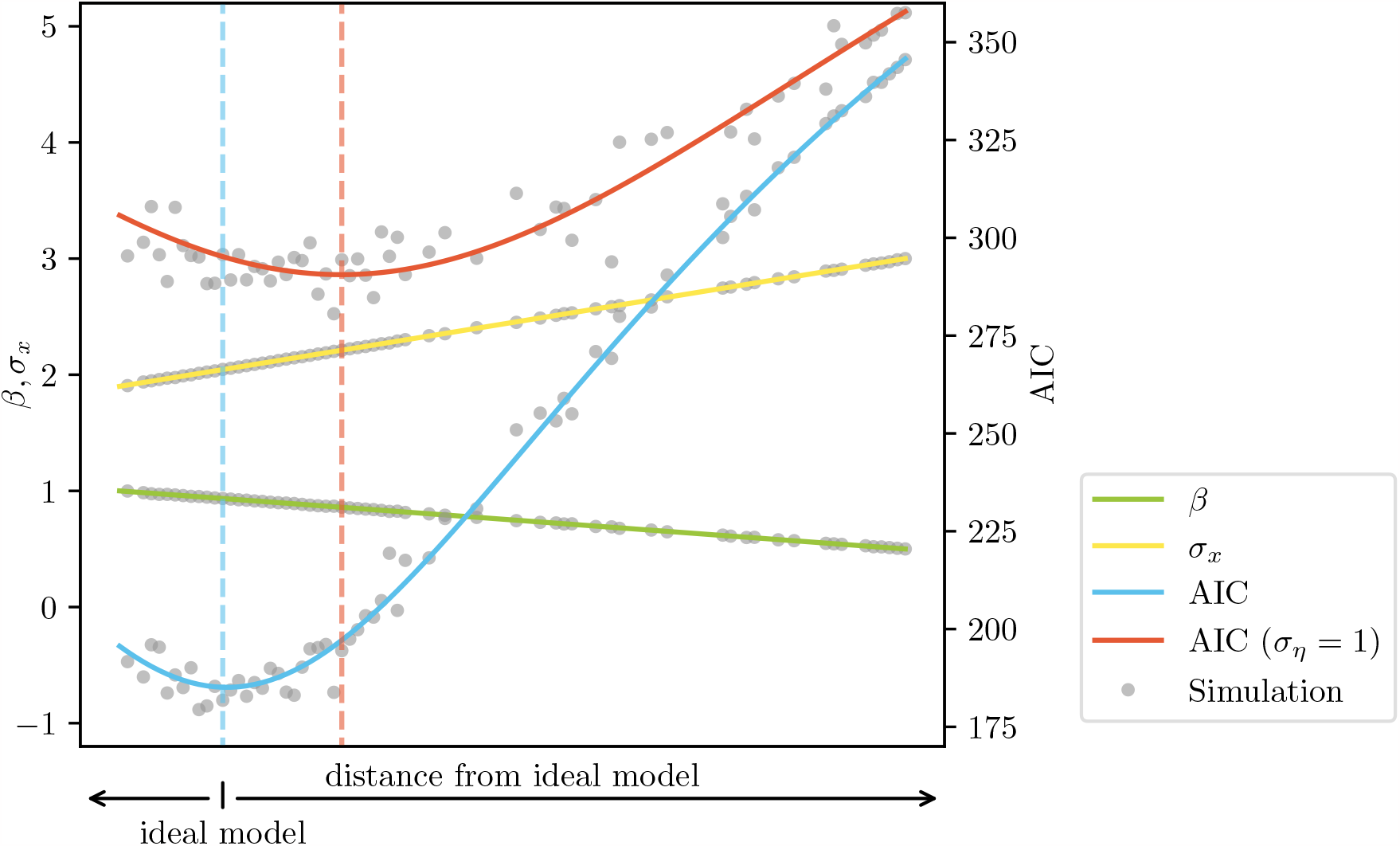
Visualisation of the ‘edges and echos’ problem. Simulating a linear decrease in effect size (*β*, green) and a linear increase of predictor variance (*σ*_*x*_, yellow) for models autocorrelated with the ideal model results in a shift of the observed best predictor (dashed red line) when using AIC of the predictor variable with added predictor error (*σ*_*η*_; shown in red) compared to the model chosen using the AIC value of the unperturbed predictor (i.e. the window that ‘should’ be chosen; shown in blue) Grey background dots show values obtained from simulation, used to validate the arithmetically derived curves.

### 5.2 Are there solutions to this problem?

Whilst it is near-impossible to quantify the extent of the predictor error problem in the extant literature, we believe it is likely that predictor error is affecting ecological model selection in a nontrivial way. As such, suggestions regarding how to mitigate against predictor error are of substantial value.

In instances where model selection is conducted using the same number of parameters in each model (i.e. where r^2^ will not be inflated by the inclusion of multiple predictors), then r^2^ possibly represents an unbiased and reliable estimate of goodness-of-fit. This statistic is, however, not calculable for all types of statistical model.

As an alternative, error-in-variable models present perhaps the most complete solution at this point. Such models seek to parameterise a causal relationship whilst accounting for an *a priori* determined amount of error in the x-axis, and in turn such models can (and have) been expanded to allow for the error-free estimation of likelihood-based goodness-of-fit metrics (e.g. AIC; Liang and Li, 2009). If error is easily estimated - for instances in systems where there is a known, causal link and measurement error can be derived - such methods might work well. Our example of wood anemone flowering date for instance could be used as a reference to estimate measurement error in the instance of proxy error, in cases where a biological cause is known, but exact measurements of the true predictor are unavailable (see Figure 3). On the other hand, in noisy systems where error is hard to quantify - for example when variables are remotely sensed and causal links are unknown - then estimating the error coefficient is far harder and as such using error-in-variable models becomes unfeasible. This, we submit, is inherently problematic. As a consequence, we suggest to consider the results of model selection carefully when error in the predictor is large or unknown.

### 5.3 Conclusion

Irrespective of the precise mechanism used to attempt to correct estimates of goodness-of-fit, we believe that our results highlight the importance of understanding the assumptions underlying statistical modeling *a priori*. More generally, we suggest that errors such as those caused by a quantity that is extremely difficult to estimate accurately, makes the effects of predictor error on model selection inherently unpredictable. This is particularly true when considering that in this study, we have only parameterised this problem for a simple linear model, and the effect of predictor error on likelihood is likely to be model-dependent. Indeed, given the complexity of statistical models used in modern ecology, an entire journal issue could likely be filled with the myriad facets of the problem we highlight here when translated to mixed-effects models, generalised linear models and other complex yet widely-used techniques. As such, we submit that the problem of predictor error warrants substantial research effort going forward.

## Supporting information

Derivations of main equations

## Software availability

All code is available at 10.5281/zenodo.10256474.

## Acknowledgements

Thanks go to Maria Moiron and Corinna Langebrake for extended discussions about this work. We also thank the Deutsche Forschungsgemeinschaft for financial support (SFB 1372 Magnetoreception and Navigation in Vertebrates, no. 395940726 awarded to ML employing GM).

## Box 1

**Constraints on variance and gradient in a model selection framework**

In model selection, the response variable (*y*) is always the same when comparing different models. This imposes certain restrictions on all other meta-variables in the system: x-variance 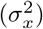, y-variance 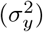, gradient (*β*), and residual-variance 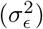. In particular, we can show that when comparing linear models, these meta-variables have to always fulfil the following equation (see SI derivation 1):

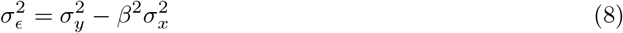

As the response variable is the same between different models, 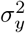 is constant. The x-variance can be directly measured or set to 1 if normalization of predictors is performed, which results in a relationship between the residual-variance and the gradient that is highly predictable (see Figure 1).

