## Supplementary material for "The unexpected consequences of predictor error in ecological model selection": Derivations of main equations

### 1 Derivations

#### 1.1 Definitions and minor results

As a basis, we consider the simple linear model including predictor-error defined as

$$\begin{aligned} y &= \beta x + \alpha + \varepsilon \\ w &= x + \eta \end{aligned} \tag{1}$$

With response $y$, predictor $x$, effect size $\beta$, intercept $\alpha$, residuals $\varepsilon$, predictor-error $\eta$ and predictor including error $w$.

In the following equations, we denote the mean or expectation of a random variable $x$ as $\bar{x} = E(x)$ and the variance as $\sigma_x^2 = V(x)$.

For our error-variables $\varepsilon$ and $\eta$, the mean is defined as 0:

$$\bar{\varepsilon} = \bar{\eta} = 0 \tag{2}$$

For our main derivations, we are mainly using the following properties of the expecation of random varaibles $x$ and $y$ and constant $c$:

$$E(x + y) = \bar{x} + \bar{y} \tag{3}$$

$$E(cx) = c \cdot E(x) \tag{4}$$

$$E(xy) = \bar{x}\bar{y} + \mathrm{Cov}(x, y) \tag{5}$$

$$E(x^2) = \bar{x}^2 + \sigma_x \tag{6}$$

From this we can already derive specific cases when variables interact with error-variables ($\varepsilon$ or $\eta$):

$$E(x\varepsilon) = \bar{x}\bar{\varepsilon} + \mathrm{Cov}(x, \varepsilon) = \bar{x} \cdot 0 + 0 = 0 \tag{7}$$

$$E(\varepsilon^2) = \bar{\varepsilon}^2 + \sigma_\varepsilon^2 = 0 + \sigma_\varepsilon^2 = \sigma_\varepsilon^2 \tag{8}$$

We will also use the well known formalization of regression dilution, where the estimated gradient ($\hat{\beta}$) is biased compared to the true gradient ($\beta$) as follows:

$$\hat{\beta} = \frac{\beta\sigma_x^2}{\sigma_x^2 + \sigma_\eta^2} \tag{9}$$

Additionally, we need a couple of small results for our main derivation, starting with the estimated intercept ($\hat{\alpha}$):

$$\begin{aligned} \hat{\alpha} &= \bar{y} - \hat{\beta}\bar{w} \\ &= \bar{y} - \hat{\beta}E(x + \eta) \\ &= \bar{y} - \hat{\beta}\bar{x} + \bar{\eta} \\ &= \bar{y} - \hat{\beta}\bar{x} \end{aligned} \tag{10}$$

In the case of a linear regression as defined in Equation 1, we can derive the covariance between variables $x$ and $y$:

$$
\begin{aligned}
\mathrm{Cov}(x,y) &= E((x-\bar{x})(y-\bar{y})) \\
&= E((x-\bar{x})(\beta x+\alpha+\varepsilon-(\beta\bar{x}+\alpha))) \\
&= E((x-\bar{x})(\beta x+\varepsilon-\beta\bar{x})) \\
&= E(\beta x^2-2\beta\bar{x}x+x\varepsilon+\beta\bar{x}^2-\bar{x}\varepsilon) \\
&= E(\beta x^2)-E(2\beta\bar{x}x)+E(x\varepsilon)+E(\beta\bar{x}^2)-E(\bar{x}\varepsilon) \\
&= \beta(\bar{x}^2+\sigma_x^2)-2\beta\bar{x}^2+\beta\bar{x}^2 \\
&= \beta\sigma_x^2
\end{aligned} \tag{11}
$$

#### 1.2 Derivation 1

With this, for a simple linear regression, we can determine how the effect size, residual variance, predictor variance and response variance interact:

$$
\begin{aligned}
\sigma_\varepsilon^2 &= E\Big((y-\beta x-\alpha)^2\Big) \\
&= E(y^2-2\beta xy-2y\alpha+\beta^2x^2+2\beta x\alpha+\alpha^2) \\
&= E(y^2)-E(2\beta xy)-E(2y\alpha)+E(\beta^2x^2)+E(2\beta x\alpha)+E(\alpha^2) \\
&= \bar{y}^2+\sigma_y^2-2\beta\bar{x}\bar{y}-2\beta\ \mathrm{Cov}(x,y)-2\bar{y}\alpha+\beta^2\bar{x}^2+\beta^2\sigma_x^2+2\beta\bar{x}\alpha+\alpha^2 \\
&= \bar{y}^2+\sigma_y^2-2\beta\bar{x}\bar{y}-2\beta^2\sigma_x^2-2\bar{y}\alpha+\beta^2\bar{x}^2+\beta^2\sigma_x^2+2\beta\bar{x}\alpha+\alpha^2 \\
&= (\beta\bar{x}+\alpha)^2+\sigma_y^2-2\beta\bar{x}(\beta\bar{x}+\alpha)-\beta^2\sigma_x^2-2\alpha(\beta\bar{x}+\alpha)+\beta^2\bar{x}^2+2\beta\bar{x}\alpha+\alpha^2 \\
&= \beta^2\bar{x}^2+2\beta\bar{x}\alpha+\alpha^2+\sigma_y^2-2\beta^2\bar{x}^2-2\beta\bar{x}\alpha-\beta^2\sigma_x^2-2\alpha\beta\bar{x}-2\alpha^2+\beta^2\bar{x}^2+2\beta\bar{x}\alpha+\alpha^2 \\
&= \sigma_y^2-\beta^2\sigma_x^2
\end{aligned}
$$

#### 1.3 Derivation 2

We can now also derive the effect of predictor error on the residual variance:

$$
\begin{aligned}
\hat{\sigma}_\varepsilon &= E((y_i - \hat{\beta} w_i - \hat{\alpha})^2) \\
&= E((\beta x_i + \alpha + \varepsilon - \hat{\beta}(x_i + \eta) - \bar{y} + \hat{\beta}\bar{x})^2) \\
&= E((\beta x_i + \alpha + \varepsilon - \hat{\beta} x_i - \hat{\beta}\eta - \beta\bar{x} - \alpha - \bar{\varepsilon} + \hat{\beta}\bar{x})^2) \\
&= E((\beta x_i + \varepsilon - \hat{\beta} x_i - \hat{\beta}\eta - \beta\bar{x} + \hat{\beta}\bar{x})^2) \\
&= E\Big(\beta^2 x_i^2 + 2\beta x_i \varepsilon - 2\beta\hat{\beta} x_i^2 - 2\beta\hat{\beta} x_i \eta - 2\beta^2 \bar{x} x_i + 4\beta\hat{\beta}\bar{x} x_i \\
&\quad + \varepsilon^2 - 2\hat{\beta} x_i \varepsilon - 2\hat{\beta}\eta\varepsilon - 2\beta\bar{x}\varepsilon + 2\hat{\beta}\bar{x}\varepsilon \\
&\quad + \hat{\beta}^2 x_i^2 + 2\hat{\beta}^2 x_i \eta - 2\hat{\beta}^2 \bar{x} x_i \\
&\quad + \hat{\beta}^2 \eta^2 + 2\beta\hat{\beta}\bar{x}\eta - 2\hat{\beta}^2 \bar{x}\eta \\
&\quad + \beta^2 \bar{x}^2 - 2\beta\hat{\beta}\bar{x}^2 \\
&\quad + \hat{\beta}^2 \bar{x}^2\Big) \\
&= E(\beta^2 x_i^2) + E(2\beta x_i \varepsilon) - E\Big(2\beta\hat{\beta} x_i^2\Big) - E\Big(2\beta\hat{\beta} x_i \eta\Big) - E(2\beta^2 \bar{x} x_i) + E\Big(4\beta\hat{\beta}\bar{x} x_i\Big) \\
&\quad + E(\varepsilon^2) - E\Big(2\hat{\beta} x_i \varepsilon\Big) - E\Big(2\hat{\beta}\eta\varepsilon\Big) - E(2\beta\bar{x}\varepsilon) + E\Big(2\hat{\beta}\bar{x}\varepsilon\Big) \\
&\quad + E\Big(\hat{\beta}^2 x_i^2\Big) + E\Big(2\hat{\beta}^2 x_i \eta\Big) - E\Big(2\hat{\beta}^2 \bar{x} x_i\Big) \\
&\quad + E\Big(\hat{\beta}^2 \eta^2\Big) + E\Big(2\beta\hat{\beta}\bar{x}\eta\Big) - E\Big(2\hat{\beta}^2 \bar{x}\eta\Big) \\
&\quad + E(\beta^2 \bar{x}^2) - E\Big(2\beta\hat{\beta}\bar{x}^2\Big) \\
&\quad + E\Big(\hat{\beta}^2 \bar{x}^2\Big) \\
&= \beta^2 \bar{x}^2 + \beta^2 \sigma_x^2 - 2\beta\hat{\beta}\bar{x}^2 - 2\beta\hat{\beta}\sigma_x^2 - 2\beta^2 \bar{x}^2 + 4\beta\hat{\beta}\bar{x}^2 \\
&\quad + \sigma_\varepsilon^2 + \hat{\beta}^2 \bar{x}^2 + \hat{\beta}^2 \sigma_x^2 - 2\hat{\beta}^2 \bar{x}^2 + \hat{\beta}^2 \sigma_\eta^2 + \beta^2 \bar{x}^2 - 2\beta\hat{\beta}\bar{x}^2 + \hat{\beta}^2 \bar{x}^2 \\
&= \beta^2 \sigma_x^2 - 2\beta\hat{\beta}\sigma_x^2 + \sigma_\varepsilon^2 + \hat{\beta}^2 \sigma_x^2 + \hat{\beta}^2 \sigma_\eta^2 \\
&= \beta^2 \sigma_x^2 - \frac{2\beta^2 \sigma_x^4}{\sigma_x^2 + \sigma_\eta^2} + \frac{\beta^2 \sigma_x^6}{\left(\sigma_x^2 + \sigma_\eta^2\right)^2} + \frac{\beta^2 \sigma_x^4 \sigma_\eta^2}{\left(\sigma_x^2 + \sigma_\eta^2\right)^2} + \sigma_\varepsilon^2 \\
&= \frac{\beta^2 \sigma_x^6 + 2\beta^2 \sigma_x^4 \sigma_\eta^2 + \beta^2 \sigma_x^2 \sigma_\eta^4 - 2\beta^2 \sigma_x^6 - 2\beta^2 \sigma_x^4 \sigma_\eta^2 + \beta^2 \sigma_x^6 + \beta^2 \sigma_x^4 \sigma_\eta^2}{\left(\sigma_x^2 + \sigma_\eta^2\right)^2} + \sigma_\varepsilon^2 \\
&= \frac{\beta^2 \sigma_x^4 \sigma_\eta^2 + \beta^2 \sigma_x^2 \sigma_\eta^4}{\left(\sigma_x^2 + \sigma_\eta^2\right)^2} + \sigma_\varepsilon^2 \\
&= \frac{\beta^2 \sigma_x^2 \sigma_\eta^2 \left(\sigma_x^2 + \sigma_\eta^2\right)}{\left(\sigma_x^2 + \sigma_\eta^2\right)^2} + \sigma_\varepsilon^2 \\
&= \frac{\beta^2 \sigma_x^2 \sigma_\eta^2}{\sigma_x^2 + \sigma_\eta^2} + \sigma_\varepsilon^2
\end{aligned}
$$

#### 1.4 Derivation 3

From derivations 1 and 2, we take into account the co-variance of the meta variables into the effect of predictor error on the residual variance:

$$
\begin{aligned}
\hat{\sigma}_\varepsilon &= \frac{\beta^2\sigma_x^2\sigma_\eta^2}{\sigma_x^2+\sigma_\eta^2}+\sigma_\varepsilon^2 \\
&= \frac{\frac{\sigma_y^2-\sigma_\varepsilon^2}{\sigma_x^2}\sigma_x^2\sigma_\eta^2}{\sigma_x^2+\sigma_\eta^2}+\sigma_\varepsilon^2 \\
&= \frac{\sigma_y^2\sigma_\eta^2-\sigma_\varepsilon^2\sigma_\eta^2}{\sigma_x^2+\sigma_\eta^2}+\sigma_\varepsilon^2 \\
&= \frac{\sigma_y^2\sigma_\eta^2-\sigma_\varepsilon^2\sigma_\eta^2+\sigma_\varepsilon^2\sigma_x^2+\sigma_\varepsilon^2\sigma_\eta^2}{\sigma_x^2+\sigma_\eta^2} \\
&= \frac{\sigma_y^2\sigma_\eta^2+\sigma_\varepsilon^2\sigma_x^2}{\sigma_x^2+\sigma_\eta^2}
\end{aligned}
$$
